# Fast inhibition slows and desynchronizes mouse auditory efferent neuron activity

**DOI:** 10.1101/2023.12.21.572886

**Authors:** Matthew Fischl, Alia Pederson, Rebecca Voglewede, Hui Cheng, Jordan Drew, Lester Torres Cadenas, Catherine J.C. Weisz

## Abstract

The encoding of acoustic stimuli requires precise neuron timing. Auditory neurons in the cochlear nucleus (CN) and brainstem are well-suited for accurate analysis of fast acoustic signals, given their physiological specializations of fast membrane time constants, fast axonal conduction, and reliable synaptic transmission. The medial olivocochlear (MOC) neurons that provide efferent inhibition of the cochlea reside in the ventral brainstem and participate in these fast neural circuits. However, their modulation of cochlear function occurs over time scales of a slower nature. This suggests the presence of mechanisms that restrict MOC inhibition of cochlear function. To determine how monaural excitatory and inhibitory synaptic inputs integrate to affect the timing of MOC neuron activity, we developed a novel in vitro slice preparation (‘wedge-slice’). The wedge-slice maintains the ascending auditory nerve root, the entire CN and projecting axons, while preserving the ability to perform visually guided patch-clamp electrophysiology recordings from genetically identified MOC neurons. The ‘in vivo-like’ timing of the wedge-slice demonstrates that the inhibitory pathway accelerates relative to the excitatory pathway when the ascending circuit is intact, and the CN portion of the inhibitory circuit is precise enough to compensate for reduced precision in later synapses. When combined with machine learning PSC analysis and computational modeling, we demonstrate a larger suppression of MOC neuron activity when the inhibition occurs with in vivo-like timing. This delay of MOC activity may ensure that the MOC system is only engaged by sustained background sounds, preventing a maladaptive hyper-suppression of cochlear activity.

**Significance Statement:** Auditory brainstem neurons are specialized for speed and fidelity to encode rapid features of sound. Extremely fast inhibition contributes to precise brainstem sound encoding. This circuit also projects to medial olivocochlear (MOC) efferent neurons that suppress cochlear function to enhance detection of signals in background sound. Using a novel brain slice preparation with intact ascending circuitry, we show that inhibition of MOC neurons can also be extremely fast, with the speed of the circuit localized to the cochlear nucleus. In contrast with the enhancement of precision afforded by fast inhibition in other brainstem auditory circuits, inhibition to MOC neurons instead has a variable onset that delays and desynchronizes activity, thus reducing precision for a slow, sustained response to background sounds.

## Introduction

The encoding of acoustic stimuli is enhanced and sharpened by active cochlear processes termed ‘cochlear amplification’, including electromotility of outer hair cells (OHC)^1–4^. Medial olivocochlear (MOC) neurons provide efferent feedback to the OHCs to inhibit this electromotility^5–10^. The subsequent suppression of sound-evoked basilar membrane vibrations improves detection of salient sounds in background noise, provides protection from noise-evoked damage, and may contribute to auditory attention^11–16^. While the impact of the cholinergic MOC synapses onto OHCs is well-characterized at the synaptic output level, the synaptic inputs onto MOC neurons in the ventral nucleus of the trapezoid body (VNTB) within the superior olivary complex (SOC) are incompletely characterized. Recent work in positively identified MOC neurons in brain slices from transgenic mice has demonstrated that cochlear nucleus (CN) T-stellate cells provide ascending afferent excitation, along with descending excitation from the inferior colliculus (IC)^17^. MOC neurons also receive afferent inhibition from the medial nucleus of the trapezoid body (MNTB) which can delay spontaneous APs in vitro^18^, and may prevent MOC neuron suppression of rapidly changing auditory stimuli^19^. MOC neuron properties allow for high frequency APs (>300 Hz) in vitro^17,20,21^. However, in vivo, MOC neurons rarely fire APs faster than 100 Hz even with high intensity sound stimulation^22–25^, and have variable latencies to APs^23,24,26^. These results indicate that MOC neurons do not operate as a simple reflex that is immediately active in response to excitatory drive from the CN. Rather, a combination of excitatory, inhibitory, and modulatory inputs may shape MOC activity in changing sound conditions.

The ascending auditory pathways from the cochlea through the brainstem have electrical and morphological specializations for temporal precision and fidelity. In particular, the three-neuron pathway (AN–GBC–MNTB) that provides the primary inhibitory synapses throughout the SOC are among the fastest, most powerful, high-fidelity circuits in the brain^27–30^, although mice have fewer GBC specializations compared to other species^31^. It is unknown whether the MNTB– GBC pathway projections to MOC neurons are also exceptionally fast, or how the timing of inhibition affects MOC neuron function.

Brain slice preparations used for synaptic physiology studies allow extensive pharmacological manipulations of circuitry, but often sever long-distance or circuitous projections, thus losing the cellular interactions that govern the timing, strength, and plasticity of incoming neuron pathways. Therefore, we developed a novel asymmetric slice preparation, the ‘wedge-slice’, that maintains sound-evoked monaural ascending circuitry^32^. This preparation enables investigation into how integration of ascending excitation and inhibition affects MOC neuron function. On one side, the slice is thin enough to allow the light penetration needed for visualization during typical whole-cell patch clamp electrophysiology experiments from MOC neurons. On the other (contralateral) side, where primary excitatory and inhibitory auditory inputs originate, the slice is thicker. This maintains the auditory nerve root as it enters the CN, the full circuitry of the CN, and the primary excitatory (direct) and inhibitory (via MNTB) axon projections from the CN to the VNTB where MOC neuron recordings are performed.

We compared the synaptic inputs to MOC neurons when stimulated at the midline (MdL) to bypass intrinsic CN circuitry vs stimulating the auditory nerve (AN) root to engage the full complement of CN circuits. We found that GBC-MNTB projections to MOC neurons are exceptionally fast and temporally precise, similar to projections to other auditory nuclei. Both the speed and precision of the circuit can be attributed to GBC neurons compensating for reduced precision in later synapses and highlighting the ‘coincidence detector’ function of GBCs^33–35^. A computational model of MOC neurons demonstrated that the enhanced speed of the AN–GBC– MNTB–MOC inhibitory pathway as measured during in vivo-like wedge-slice conditions has a larger effect on suppressing activity in MOC neurons compared to with simple stimulation (MdL-stimulation). This increased speed of inhibition delayed AP onset in MOC neurons, while variability of inhibitory timing across the population of MOC neurons results in stochastic activity that may provide smooth inhibition of cochlear function.

## Results

### Midline stimulation in wedge-slice preparations evokes mixed excitatory and inhibitory post-synaptic currents (PSCs)

Ascending axons conveying acoustic information from the CN throughout the SOC traverse near the ventral surface of the brainstem. This includes axons projecting to MOC neurons in the VNTB that provide sound-evoked excitation, primarily T-stellate and possibly small cell cap (SCC) neurons^17,36–39^. This also includes axons of GBCs which provide sound-evoked inhibition via intervening MNTB neurons^18^. To investigate the convergence of these pathways, whole cell patch-clamp recordings were performed in voltage-clamp from positively identified MOC neurons in brainstem slices from postnatal day fourteen through nineteen (P14-P19) ChAT-IRES-Cre; tdTomato mice of both sexes. We used a combination of uniformly thick (300 micron) slices cut at ∼15 degrees off the coronal plane^18^ and wedge-slices^32^ (**Figure 1A**) while electrically stimulating close to the ventral surface of the slice near the midline (MdL) to evoke neurotransmitter release from pre-synaptic axons. These axons likely originated in the CN contralateral to the MOC neuron. With this technique, both excitatory and inhibitory circuits originating at the contralateral CN can be simultaneously stimulated. MdL-stimulation in this location is expected to directly activate T-stellate cell axons resulting in mono-synaptically evoked, short latency excitatory post-synaptic currents (Midline-evoked EPSCs: MdL-EPSCs). Additionally, stimulation of GBC axons accessed from the same location results in inhibitory post-synaptic currents (Midline-evoked IPSCs: MdL-IPSCs) evoked via the di-synaptic GBC–MNTB-MOC pathway that inhibits MOC neurons^18^. Other as-yet uncharacterized synaptic inputs to MOC neurons may also be activated with this technique. Consistent with activation of multiple classes of pre-synaptic axons, MdL-stimulation resulted in multi-component PSCs in 13/21 recordings (**Figure 1B**). PSCs occurred with a short latency from stimulation (1.85±0.53 ms, n=557 PSCs in 21 neurons; a subset of PSC latency data was previously published^32^).

**Figure 1.**
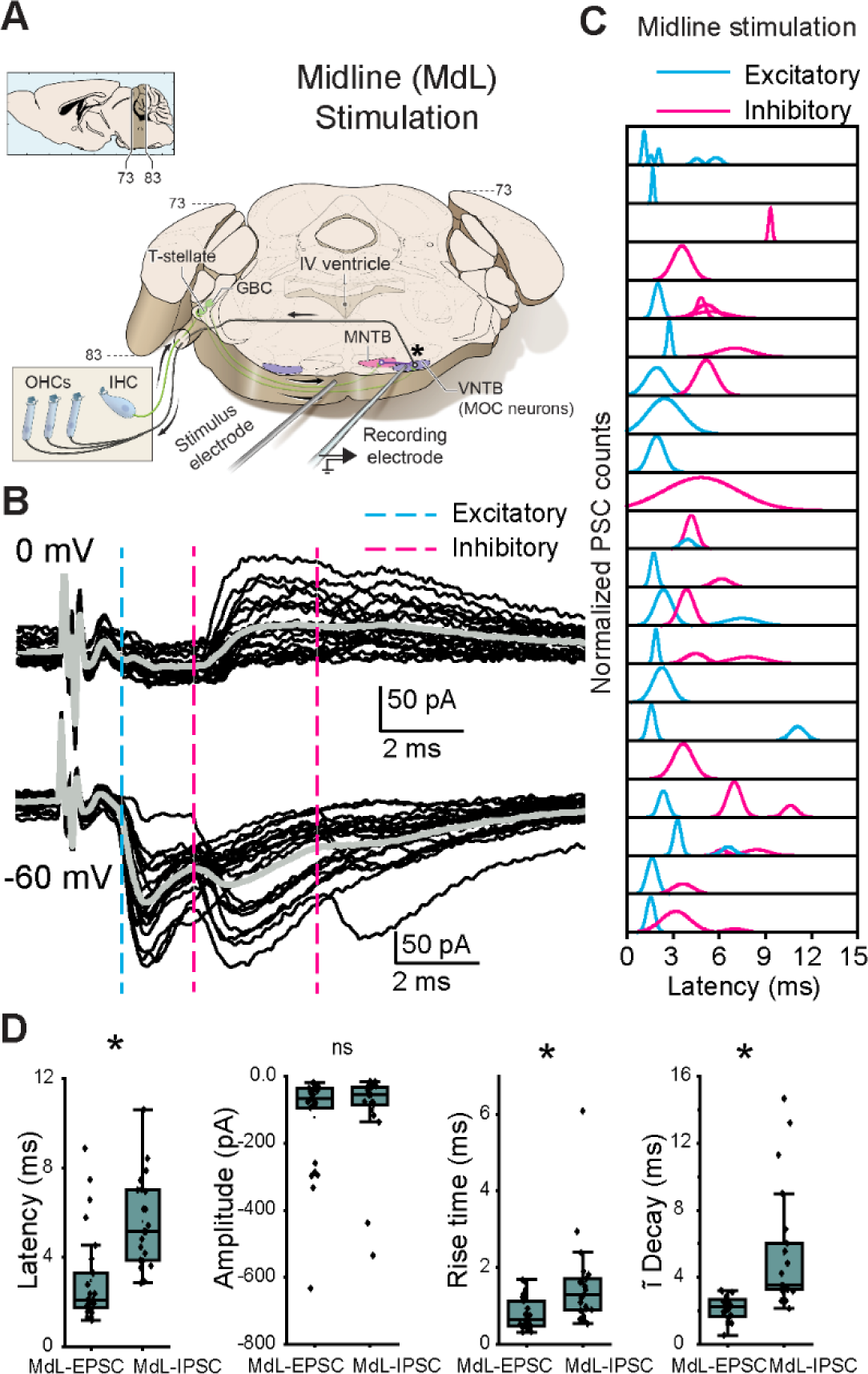
Stimulation of axons at the midline of the ventral brainstem evokes multi-component PSCs in MOC neurons. A. Schematic of wedge-slice preparation showing monaural circuitry of MOC neurons from ascending AN axons from the cochlea into the CN, excitatory (green) T-stellate and GBC projections to the SOC, inhibitory (purple) projections from the MNTB to the VNTB containing MOC neurons, and MOC axons (black) projecting back to cochlear OHCs. Note: the cochlea is not present in the recording preparation. Upper left schematic indicates atlas coordinates of position along rostral-caudal extent of thick portion of the slice^125^. B. Voltage-clamp traces from an identified MOC neuron during MdL-stimulation (1 Hz), evoking PSCs in multiple clusters at a holding potential of 0 mV (top) and -60 mV (bottom). 20 sweeps overlaid (black), with grey line indicating the average waveform. Dashed lines indicate the approximate onset of PSCs in clusters of excitatory (blue) and inhibitory (magenta) PSCs. C. Gaussian distributions fitted to frequency histograms (normalized) of the latency to PSC onset for clusters of MdL-EPSCs (blue), and MdL-IPSCs (magenta). Each row represents PSCs recorded in a different MOC neuron. D. Comparison of parameters observed in MdL-EPSCs and MdL-IPSCs.

With the high intracellular chloride concentration used in voltage-clamp recordings to maximize detection of IPSCs (intracellular chloride concentration = 60 mM, E_Cl_ ∼-20 mV), both ‘excitatory’ and ‘inhibitory’ PSCs were inward at a membrane holding potential of -60 mV, and therefore indistinguishable based on polarity. To distinguish between MdL-EPSCs and MdL-IPSCs, in all cells the MOC neuron holding potential was alternatively set to the AMPA receptor reversal potential of 0 mV to isolate outward, chloride-mediated PSCs that can be classified as ‘inhibitory’ that are likely either GABAergic or glycinergic. In each experiment, stimulation was performed repeatedly (20-80 stimulations; **Figure 1B**), and in each sweep, PSCs were detected and analyzed for parameters of onset latency, rise time (10-90% of peak), amplitude, and time constant of decay. These parameters (excluding amplitude) were used to sort PSCs into statistically defined clusters using k-means analyses (Methods). Clusters observed at both –60 mV and 0 mV holding potentials were classified as ‘inhibitory’ MdL-IPSCs and clusters observed only at –60 mV were classified as ‘excitatory’ MdL-EPSCs (**Figure 1C**, each peak indicates a different ‘cluster’).

Following classification of PSCs as ‘excitatory’ or ‘inhibitory’ based on the presence or absence of the PSC cluster at a holding potential of 0 mV, we further characterized MdL-PSCs recorded at -60 mV. Latency to the first MdL-EPSC was shorter than the latency to the first MdL-IPSC (p=4.79E-06). MdL-EPSCs and MdL-IPSCs had similar amplitudes at -60 mV (p=0.69). MdL-EPSCs had faster kinetics than MdL-IPSCs (rise time: p=8.02E-04, time constant of decay: p=4.95E-07, **Table S1, Figure 1D**), consistent with earlier work^18^. Patterns of PSCs varied somewhat across MdL stimulation experiments. Patterns included recordings with EPSCs only (6/21), IPSCs only (4/21), a single group of EPSCs and a single group of IPSCs (5/21 recordings), and more complex patterns of EPSCs and IPSCs (6/21) (**Figure 1C**). In all recordings with both EPSCs and IPSCs (11/21), EPSCs always occurred with a shorter latency than IPSCs. This shorter latency was expected given that the MdL-EPSC pathway is mono-synaptic (T-stellate-MOC) while the MdL-IPSC pathway is di-synaptic (GBC-MNTB-MOC), consistent with an extra synapse in the MdL-IPSC pathway incurring a delay.

### Auditory nerve stimulation-evoked PSCs

The above experiments detail the relative timing of synaptic inputs to MOC neurons evoked simultaneously at the MdL from mono-synaptic excitatory and di-synaptic inhibitory inputs. The timing of synaptic inputs in above experiments with MdL-stimulation is artificial because the full complexity of ascending circuitry is not activated in this stimulation paradigm, including AN synapses onto CN neurons, intrinsic CN circuits, and the potential differential propagation of APs down the specialized CN axons. To test the integration of ascending, monaural, excitatory, and inhibitory synaptic inputs to MOC neurons with a more in vivo-like timing, PSCs were evoked in MOC neurons by stimulating the contralateral AN root in wedge-slice preparations (**Figure 2A**), which have an intact CN.

**Figure 2.**
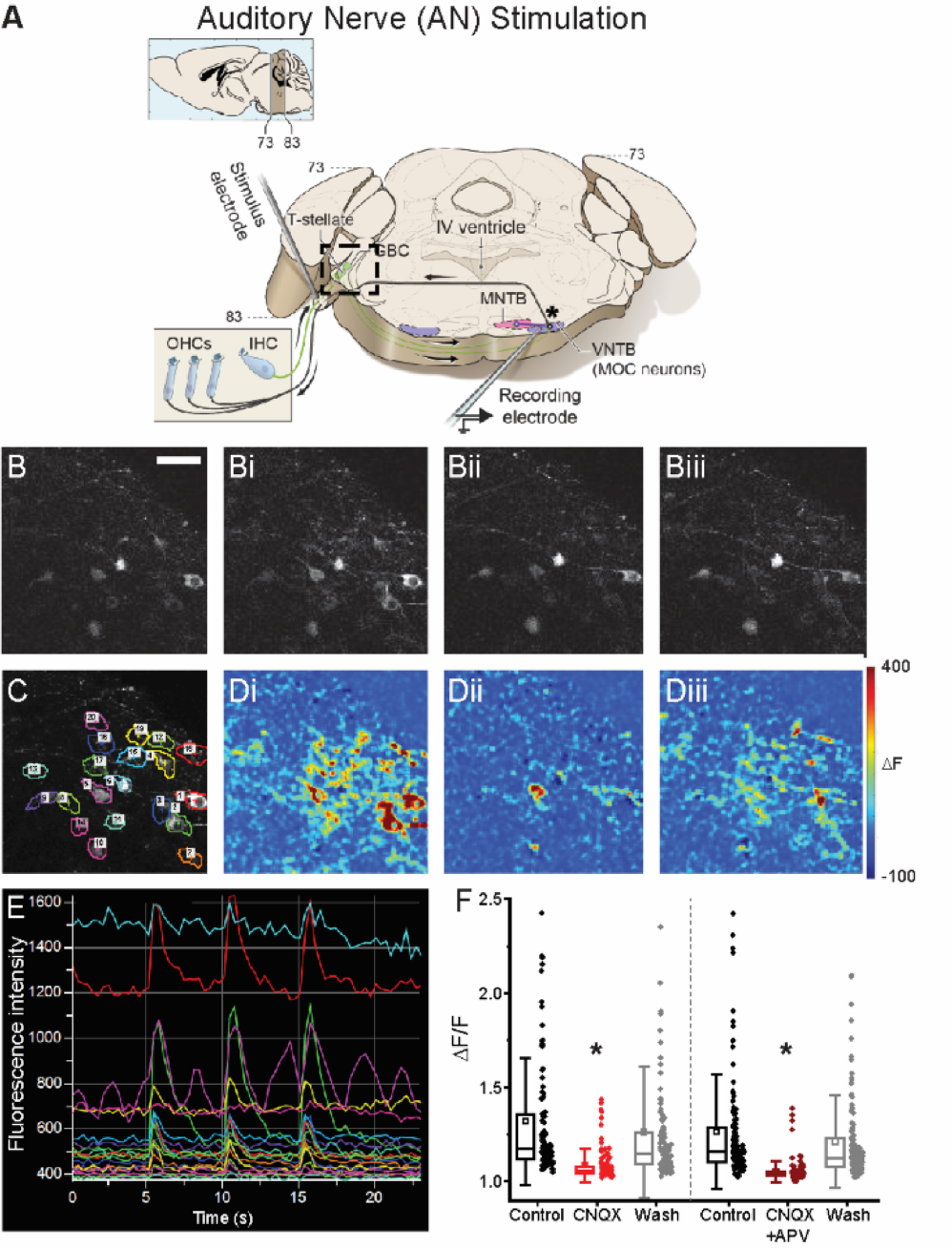
AN-stimulation in a wedge-slice evokes activity via synaptic activation of CN neurons. A Schematic of a wedge-slice, as in Figure 1, with stimulation at the auditory nerve root (AN-stimulation). Note: the cochlea is not present in the preparation. B. Calcium imaging of bushy cells in Atoh7/Math5^Cre^; GCaMP6f mice during AN-stimulation in: B. Baseline (no stimulation), Bi. control AN-stimulation at 100 Hz, 20 pulses, Bii. AN-stimulation with blockers of AMPA and NMDA receptors (5 µM CNQX, 50 µM APV) in the aCSF. Biii. AN-stimulation after wash of AMPA and NMDA receptor blockers. C. Same CN imaging region as in B, annotated with ROI’s over individual bushy cells used for fluorescence analysis. D. Heat maps showing change in fluorescence in same region as B and C during AN-stimulation, color scale indicates fluorescence change from baseline in: Di. control, Dii. AMPA and NMDA block, and Diii. wash. E. Raw fluorescence intensity changes for ROIs in C during 3 ‘bouts’ (bout: 20 pulses, 100 Hz) of AN-stimulation in control conditions. F. Quantification of fluorescence changes of bushy cells in response to AN-stimulation in control, glutamate receptor blockade (CNQX or CNQX + APV) and wash. Scale bar in B (50 μm) applies to all panels in B-D.

To ensure that electrical stimulation indeed excited CN neurons via synaptic activation, and not direct electrical activation which would bypass the strength and timing of AN–CN synapses, we performed 2-photon calcium imaging of bushy cells in wedge-slices from Atoh7/Math5^Cre^ mice^40,41^ expressing GCaMP6f^42^ in bushy cells, to measure supra-threshold activation during AN-stimulation. AN axons were stimulated in control conditions and with bath application of glutamate receptor antagonists to block synaptic transmission at AN–CN synapses. If the AN electrical activation stimulates glutamate release from AN axons to synaptically activate bushy cells, blockade of the glutamate receptors is expected to reduce the somatic calcium response. However, if AN electrical stimulation directly evokes APs in bushy cells, the calcium response would be insensitive to glutamate receptor blockers.

AN-stimulation-evoked calcium responses were quantified for 388 neurons in 14 fields of view from 8 animals (**Figure 2B,E**). Of these neurons, 227 were classified as ‘active’ in the control condition. AN-evoked calcium responses in active bushy cells were significantly reduced by bath application of the AMPA receptor antagonist CNQX (p=1.04E-20) or the combination of CNQX and the NMDA receptor antagonist APV (p=1.14E-34). Calcium responses significantly recovered upon wash of glutamate receptor blockers (CNQX recovery: p=2.74E-13; CNQX+APV recovery: p=2.54E-24). There was a small but significant difference in the ΔF/F values between the CNQX and the CNQX+APV groups (p=0.005) (**Figure 2B-D**), confirming active NMDA receptors in bushy cells^43^. We explored this difference further by calculating a % suppression for each of the neurons. CNQX suppressed the calcium signal by ∼68% while the cocktail of CNQX+APV suppressed the signal by ∼75% (p=0.004) (**Table S2, Figure 2F**). Across cells, calcium suppression with CNQX and APV was significant but not quite complete, perhaps due to incomplete penetration of antagonists into the thick wedge-slice. However, it is notable that GBCs have a particularly small somatic AP^44–46^ and an abundance of calcium permeable glutamate receptors^43^. Therefore, the relative contribution of the AP-evoked calcium signal to the total evoked calcium signal may be small, resulting in a relatively large residual calcium signal in the case of incomplete glutamate receptor block even if APs are suppressed. However, the sensitivity of calcium responses in bushy cells to glutamate receptor blockers suggests that AN-stimulation is likely activating CN neurons via synaptic, and not direct electrical excitation, maintaining the AN-CN synapses in the wedge-slice circuit for a more in vivo-like network activation.

### Multi-component synaptic responses evoked by AN-stimulation

To test how the extensive intrinsic circuitry of the intact CN and specialized axons projecting to the SOC change the relative timing of excitatory and inhibitory synaptic inputs to MOC neurons, the AN was electrically stimulated in a wedge-slice while PSCs were recorded in contralateral MOC neurons (**Figure 3A**). Electrical stimulation at the AN root evoked PSCs in 11 of 43 MOC recordings. Similar to MdL-evoked PSCs, AN-evoked PSCs at a membrane holding potential of -60 mV occurred in multi-component PSC patterns (**Figure 3B**). Overall, the latency to the first PSC was significantly longer in AN- vs MdL-evoked PSCs (AN-PSC latency: 5.51±1.00 ms, n=11 cells; MdL-PSC latency: 1.99±0.45 ms, n=21 cells, Mann-Whitney U test p=2.29E-5), consistent with an increased total number of intervening synapses causing a longer synaptic delay.

**Figure 3.**
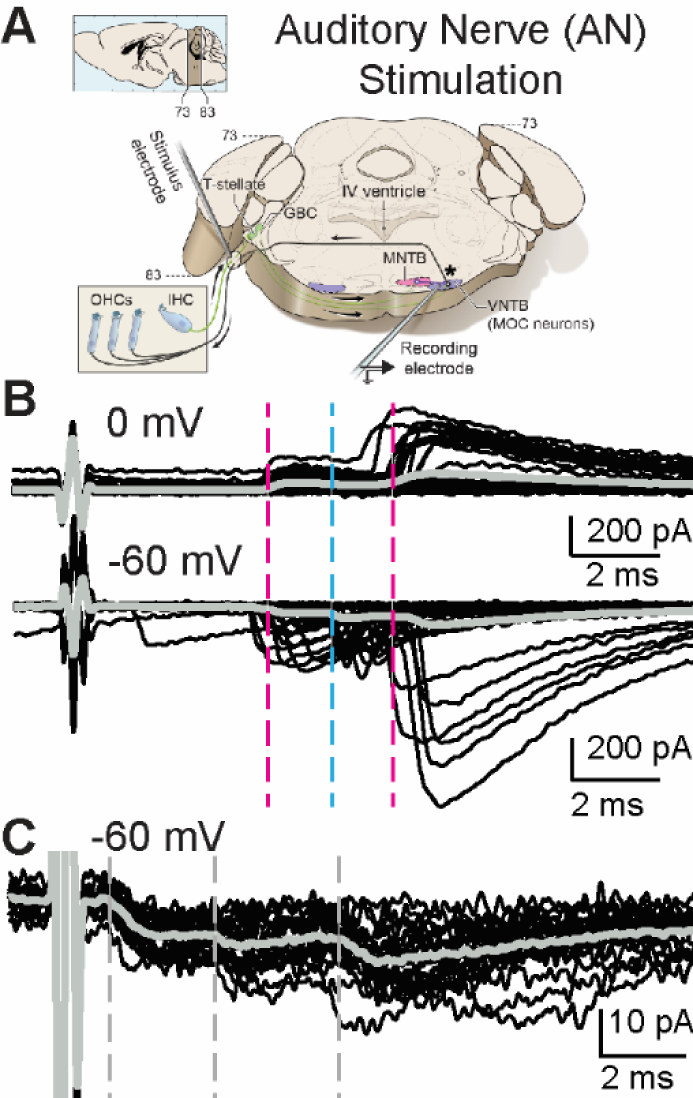
AN-stimulation evokes excitatory and inhibitory PSCs in MOC neurons. A. Schematic of the wedge-slice, indicating electrical stimulation of AN axons (AN-stimulation) projecting to the CN. B. Voltage-clamp traces from identified MOC neurons during AN-stimulation at 1Hz, 0 mV (top) and -60 mV (bottom). Black traces are 50 overlaid sweeps. Grey line indicates average trace. Dashed lines indicate approximate PSC onset latency for clusters of excitatory (blue) and inhibitory (magenta) PSCs. C. Voltage-clamp traces at -60 mV from a different MOC neuron than in (B) of PSCs evoked by AN-stimulation when the stimulus intensity is increased to levels high enough to evoke short-latency PSCs.

In two experiments, AN-stimulation protocols were repeated with high intensity electrical stimulation (>2000 μA), to cause greater current spread in the tissue and intentionally bypass the AN–CN synapse. The high stimulation intensity significantly decreased PSC latency (AN-high stim level latency: 1.65±0.1 ms, n=18 PSCs; AN-intermediate stim level latency: 4.58±0.3 ms, n=27 PSCs; n=2 neurons, Mann-Whitney U test p=1.90E-8; **Figure 3C**, **Figure 4Fi**). This indicates that with intentionally high stimulation intensity CN axons projecting to MOC neurons were directly stimulated, bypassing intrinsic circuitry of the CN. Remaining experiments used the lower stimulation intensity that synaptically engages, not electrically bypasses, the full complement of CN circuits.

**Figure 4.**
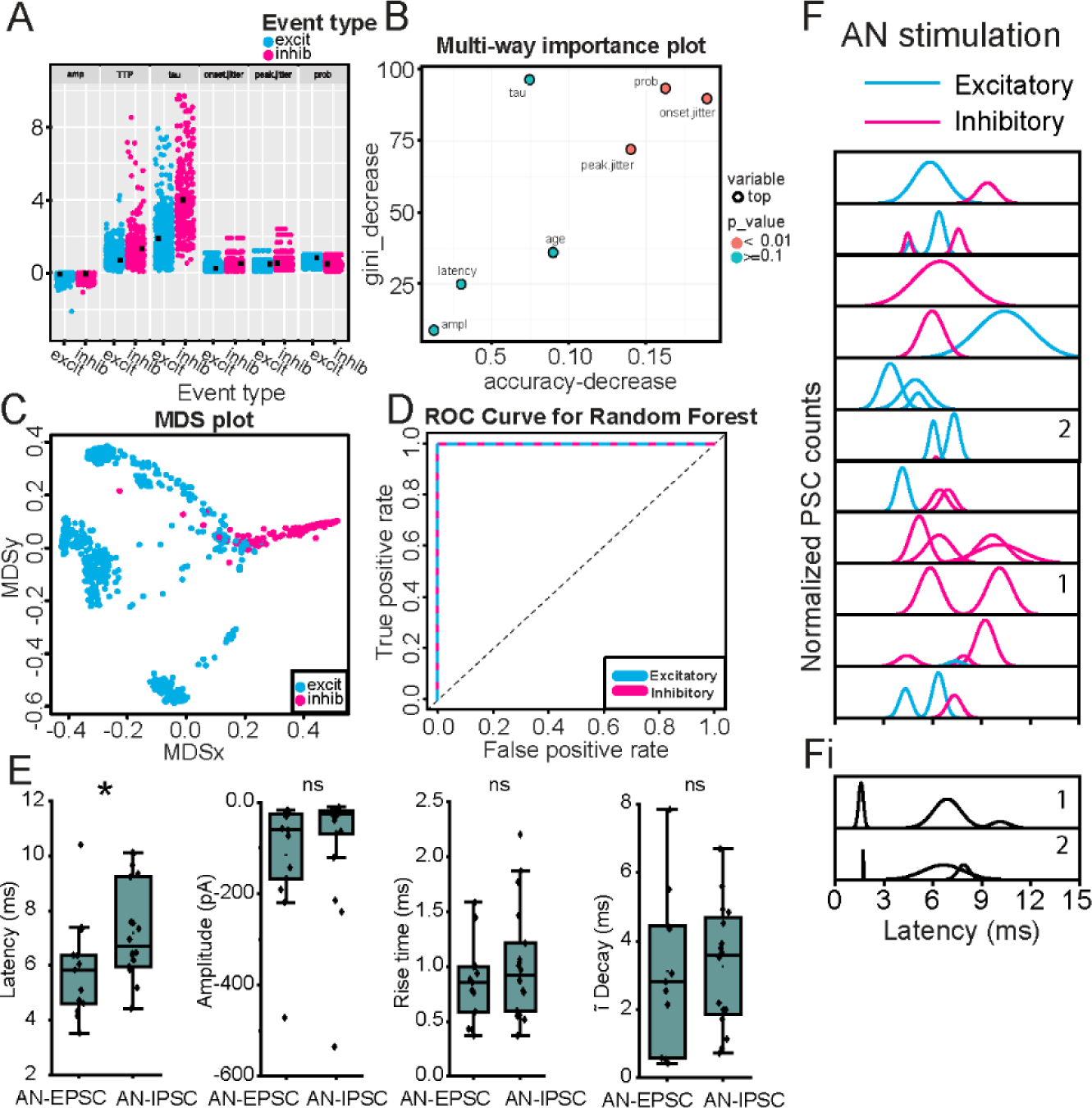
Multi-component PSCs evoked by AN-stimulation in a wedge-slice. A. Data points for each of seven variables (amp = amplitude; TTP = time to peak; tau = time constant of decay; prob = PSC probability) from MdL-EPSCs (blue) and MdL-IPSCs (magenta) used to train the RandomForest machine learning algorithm. B. Importance plot of variables used for RandomForest algorithm. C. Multidimensional Scaling (MDS) plot of proximity matrix showing separation of MdL-EPSCs and MdL-IPSC variables. D. ROC Curve for excitatory and inhibitory MdL-PSCs. The algorithm had a 99.89% classification accuracy. E. Comparison of parameters of AN-EPSCs and AN-IPSCs. F. Gaussian fits to frequency histograms (normalized) indicating the latency to clusters of AN-EPSCs (blue) and AN-IPSCs (magenta). Each row represents a different MOC neuron. Fi. Gaussian fits to frequency histograms (normalized) indicating the latency to clusters of AN-PSCs after the stimulation intensity was increased to higher levels that recruit short-latency PSCs. Numbers in panels in F and Fi indicate PSC clusters recorded from the same MOC neuron with intermediate (F) and high (Fi) stimulation intensities.

### Machine Learning Algorithm to classify excitatory and inhibitory PSCs

Analysis of synaptic integration requires classification of synaptic responses as excitatory or inhibitory. In some (4/11) experiments with AN-stimulation the MOC neuron membrane holding potential was set to 0 mV to isolate IPSCs. Similar to MdL-stimulation recordings, IPSCs occurred in clusters that aligned with a subset of the clusters recorded at -60 mV (**Figure 3B**), indicating that both EPSCs and IPSCs are evoked in MOC neurons by AN-stimulation. However, for the cells lacking 0 mV data, we were initially unable to classify PSC clusters. Therefore, we used parameters of PSCs evoked from MdL-stimulation to develop a machine learning approach to characterize EPSCs and IPSCs, and then used this algorithm of MdL-evoked PSC characteristics to classify AN-evoked PSCs as excitatory or inhibitory. To accomplish this, first we developed a RandomForest-based classification algorithm to identify EPSCs and IPSCs by training the model on identified MdL-EPSCs and MdL-IPSCs recorded at -60 mV (**Figure 4A-D**).

Next, we used the trained RandomForest algorithm to classify AN-PSCs recorded at -60 mV as excitatory or inhibitory. The 344 AN-PSCs from 11 MOC neuron recordings were clustered in the same manner as the MdL-PSCs (above), and the PSC parameters were calculated on a PSC-by-PSC basis (rise time, time constant of decay, amplitude, animal age) or a cluster-by-cluster basis (onset jitter, peak jitter, PSC probability). The AN-PSCs were then fed into the trained RandomForest algorithm, which gave the probability that each AN-PSC could be classified as AN-EPSC or AN-IPSC.

### AN-evoked synaptic responses

We then determined the effect of AN-PSCs on MOC neurons. Following classification of AN-EPSCs and AN-IPSCs responses at -60 mV using the RandomForest algorithm above, the parameters of AN-EPSCs and AN-IPSCs for all clusters were summarized to determine their effects on MOC neurons (**Table S3, Figure 4E**). AN-EPSCs and AN-IPSCs at -60 mV had similar amplitudes (p=0.14) and kinetics (rise time: p=0.35, time constant of decay p=0.67). The median AN-EPSC latency occurred with a significantly shorter latency than AN-IPSCs (p=0.04), but in two cells AN-IPSCs occurred with shorter latencies than AN-EPSCs, similar to projections to MSO neurons^28^. AN-evoked PSCs also occurred in complicated patterns that could have multiple excitatory or inhibitory clusters (**Figure 4F**, each peak indicates a different ‘cluster’). PSCs evoked by AN-stimulation were more likely to have ‘complex’ patterns with two or more clusters of either EPSCs or IPSCs (AN 8/11 vs MdL 8/21 complex clusters), interpreted here as PSCs from multiple pre-synaptic sources.

Comparison of PSCs evoked by MdL- vs AN-stimulation demonstrated a specialized function of CN circuitry, which was both intact and engaged during AN-stimulation but not MdL-stimulation. Above data (**Figure 4**) examines all AN-PSCs, but here we consider the first excitatory cluster and first inhibitory cluster to isolate our analyses to direct ascending pathways. The overall latency to the first AN-PSCs was increased compared to MdL-PSCs, as expected because AN-stimulation includes at least one more synapse (AN-CN neurons) compared to MdL-stimulation. This occurred for both EPSCs and IPSCs (MdL-EPSC vs AN-EPSC latency, p=8.98E-5; MdL-IPSC vs AN-IPSC latency, p=0.049). However, the additional circuitry added during AN-stimulation does not alter EPSC and IPSC timing to the same degree. When comparing MdL-stimulation evoked EPSCs and IPSCs, MdL-EPSCs are significantly shorter latency than MdL-IPSCs (p=4.79E-6), and MdL-IPSCs are never recorded before MdL-EPSCs in any cell. In contrast, when stimulating the AN, there is no significant difference in latency between AN-EPSCs and AN-IPSCs (p=0.30), suggesting that the CN circuitry can compensate in speed for the additional synapse present in the IPSC pathway to equalize the timing of the excitatory and inhibitory pathways (**Table S4, Figure 5A**). This is also apparent when calculating the “E-I latency difference” on a cell-by-cell basis, for which the latency to the first IPSC is subtracted from the latency to the first EPSC. MdL-stimulated EPSCs all had positive E-I latency difference values that were significantly different from zero (One-Sample Wilcoxon Signed Rank Test, p=0.0039), indicating that excitation always preceded inhibition. However, the AN-stimulation E-I latency difference was smaller than the MdL-stimulation E-I latency difference (MdL: 2.80±0.88, n=11 cells; AN: 0.14±2.87, n=7 cells, Mann-Whitney U test p=0.085), and not significantly different from zero (One-Sample Wilcoxon Signed Rank Test, p=1). Further, two AN-stimulation experiments had negative E-I latency differences, indicating that inhibition preceded excitation in these cells. These data confirm the unusual speed of the inhibitory pathway from the CN to the SOC relative to the excitatory pathway. Further, comparison of AN-stimulation and MdL-stimulation synaptic timing demonstrates that the remarkable speed of the inhibitory pathway is localized to components of the circuit added to the preparation during AN-stimulation, namely the AN synapse onto the GBCs and the GBCs themselves.

**Figure 5.**
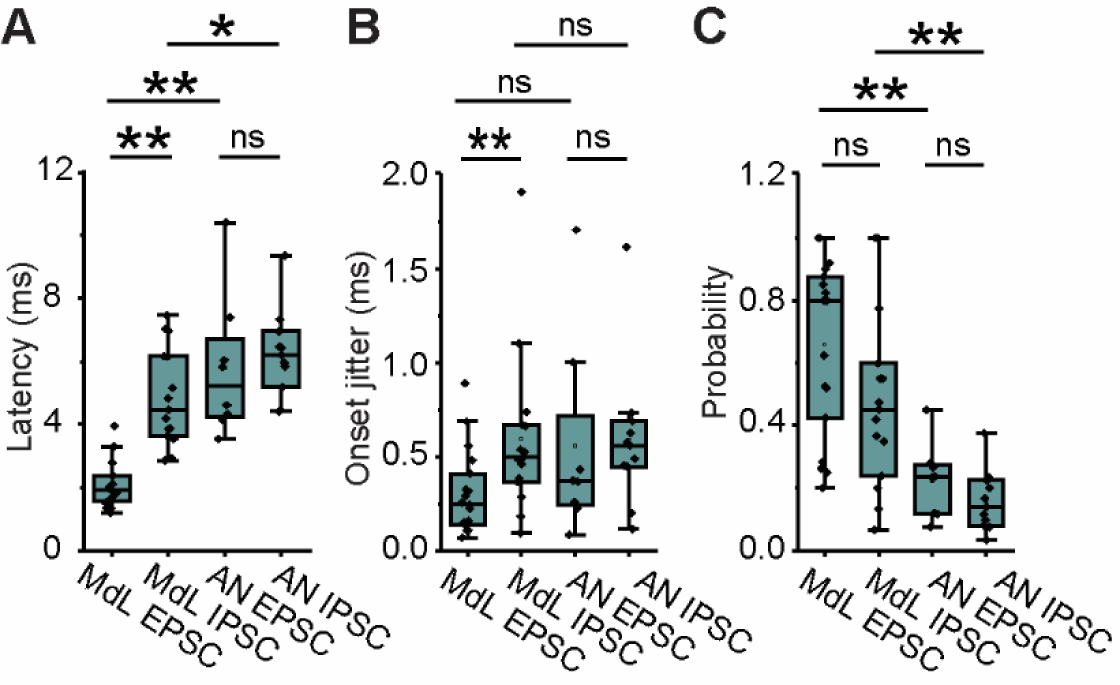
Comparison of PSCs evoked from MdL- vs AN-stimulation. A-C. Plot of A. Latency to PSC onset, B. Synaptic jitter of PSC onset, and C. Probability of recording a PSC for MdL- vs AN-EPSCs and IPSCs in the first cluster recorded in each MOC neuron.

Auditory brainstem circuits contain extremely precise neurons, so the timing and precision of synaptic inputs evoked from MdL- vs AN-stimulation experiments was investigated to determine if MOC synaptic inputs are comparably temporally precise. The onset jitter of PSCs was computed for the first EPSC cluster and first IPSC cluster per MOC neuron for both MdL- and AN-stimulation (**Figure 5B**). A simplistic view suggests that each additional synapse in a pathway would increase the overall pathway jitter, resulting in EPSCs having less jitter than IPSCs in all recording configurations. Indeed, monosynaptic T-stellate-MOC MdL-EPSCs had lower jitter than di-synaptic GBC-MNTB-MOC MdL-IPSCs (p=0.015). However, AN-EPSC jitter is not different from AN-IPSC jitter (p=0.30), The lack of difference in jitter between the excitatory and inhibitory pathways with AN-stimulation suggests that the inhibitory pathway can compensate for having an additional synapse by having enhanced precision in the CN. There was no difference in PSC probability between MdL-EPSCs and MdL-IPSCs (p=0.08), and there was also no difference in probability of AN-EPSCs and AN-IPSCs (p=0.16). However, when stimulating at the AN, both the excitatory and inhibitory pathways have a lower probability of PSCs compared to stimulating at the MdL (**Table S4, Figure 5C**). This was unexpected for the inhibitory pathway that has a high proportion of suprathreshold PSPs^34,43,45^, and unexpectedly indicates a reduced throughput at the endbulb–GBC synapse compared to stimulating the GBC–MNTB synapse alone in our experiments with MdL-stimulation.

### Summation of ascending MdL-evoked synaptic inputs drives APs in MOC neurons

The voltage-clamp experiments above demonstrate that the inhibitory pathways projecting to MOC neurons can be remarkably fast when the in vivo-like circuitry is intact. However, the impact of the timing of inhibition on MOC neuron AP activity is unknown. We tested integration under more physiological conditions of low intracellular chloride (7.2 mM, E_Cl_ = -74 mV), without intracellular QX-314, and in the current-clamp configuration. We stimulated both excitatory and inhibitory synaptic pathways simultaneously to record post-synaptic potentials (PSPs) and resulting APs in MOC neurons using the MdL-stimulation configuration because these experiments were high-throughput enough to allow pharmacological blockade of inhibitory synaptic inputs. MdL-stimulation was applied in trains to evoke PSPs (MdL-PSPs) (**Figure 6A, Ai**). Summation of MdL-PSPs evoked APs, with increased stimulation rates evoking APs with an increased probability, increased rate, and reduced latency to the first AP (Friedman ANOVA: AP probability: p=0.031, AP rate: p=0.007, number of stimulations to first AP: p=0.041, Latency to first AP: p=4.6E-4; **Figure 6A, C; Table S5**).

**Figure 6.**
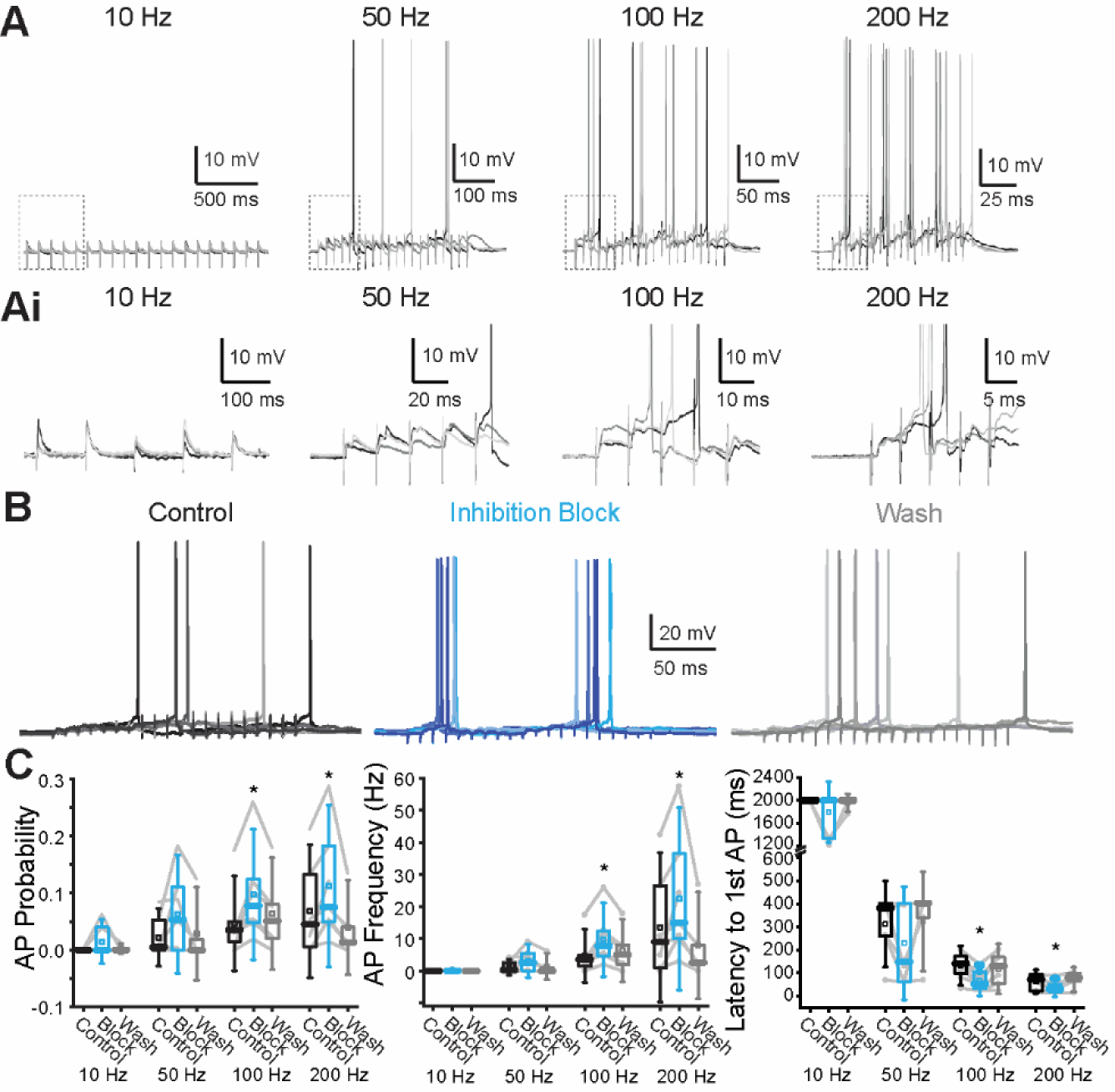
Inhibition reduces AP activity in MOC neuron recordings in response to MdL-PSPs. A. Current-clamp recordings from an MOC neuron while evoking MdL-PSPs for 20 pulses at 10, 50, 100, and 200 Hz. Three sweeps shown, overlaid, at each stimulation rate. Ai. Zoom of regions in each panel of (A) indicated by dashed box, showing the first five PSPs evoked in MdL-stimulation trains. B. 100 Hz MdL-stimulation-evoked PSPs summate to generate AP trains in control conditions (left), in the presence of 50 µM gabazine and 1 µM strychnine to block inhibitory inputs (“Inhibition block”, center), and after wash of inhibitory receptor blockers (right). 5 sweeps are overlaid per condition. C. Quantification of AP probability (left), frequency (center), and latency to first AP (right) for 5-8 neurons.

We then tested the effect of inhibition on AP rates in MOC neurons in response to MdL-stimulation by blocking inhibitory neurotransmitter receptors. In the absence of inhibitory neurotransmission, there was an increased AP probability and rate, decreased number of stimulations to the first AP, and decreased latency to the first AP for 100 and 200 Hz stimulation rates (**Figure 6B, C; Table S5**).

### Computational model of MOC neurons to assess integration of excitation and inhibition

In the MdL-stimulation experiments above, pharmacological blockade of inhibition increased APs in MOC neurons. We next asked how the effect of inhibition on MOC activity would change when excitatory and inhibitory pathways were stimulated with the in vivo-like synaptic timing that we observed in the AN-stimulation experiments. However, the short window of time for wedge-slice viability for AN-stimulation experiments made the additional pharmacological experiments necessary for testing the effect of inhibition on MOC neuron activity prohibitive. Therefore, we built a computational model of an MOC neuron to test the integration of excitation and inhibition with synaptic inputs that mimic AN-stimulation experiments. The model MOC neuron had topology generated from a published MOC neuron morphology (Figure 9^47^, **Figure 7A**), and ion channels and biophysical properties were tuned to mimic the recorded synaptic and AP activity of MOC neurons (Methods).

**Figure 7.**
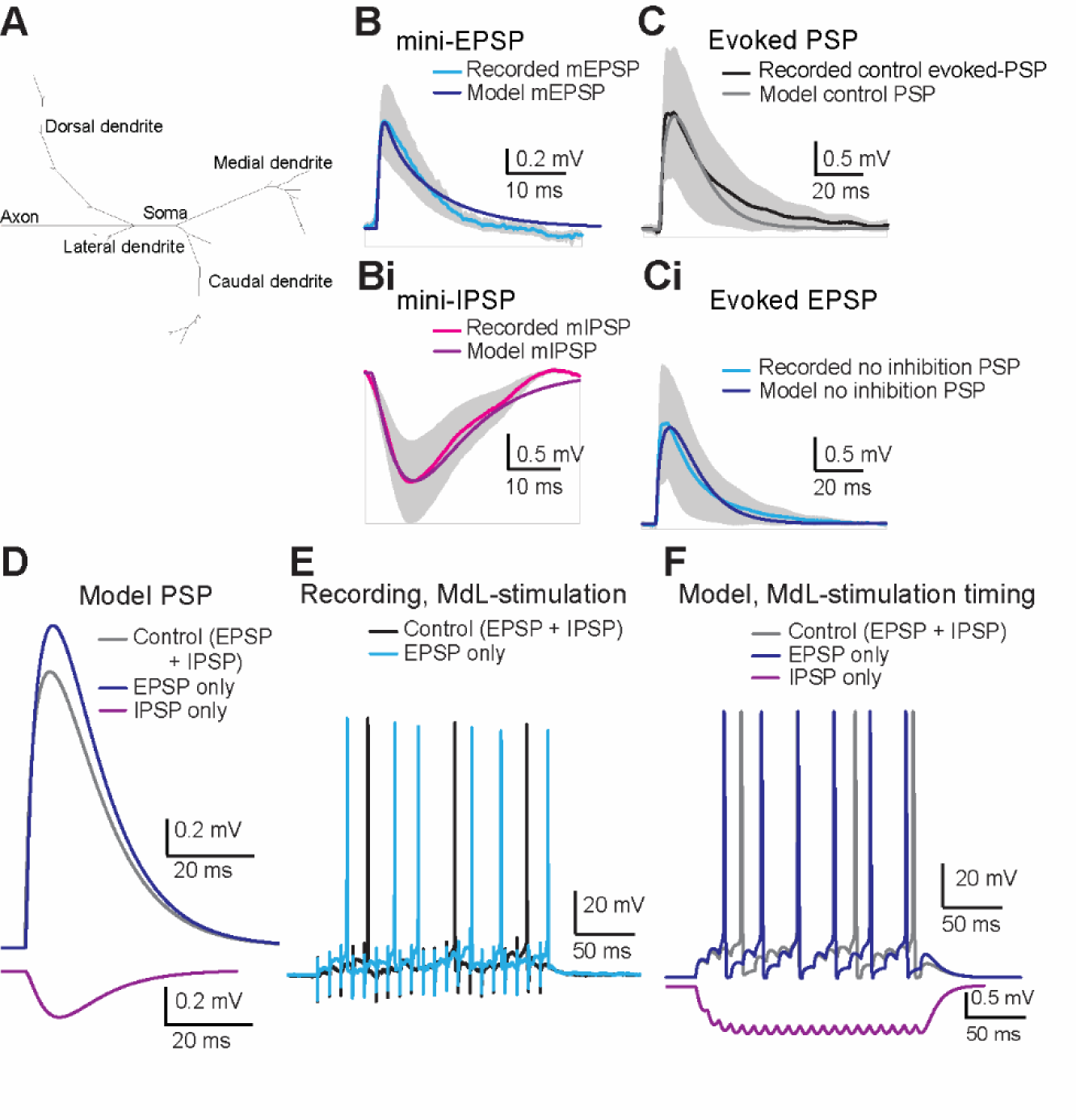
A computational MOC model replicates responses to MdL-stimulation. A. Topology of the model MOC neuron. B. Waveform of mEPSPs from MOC neuron recordings (light blue line, grey shading indicates standard deviation) with overlaid mEPSP generated from the model MOC neuron (dark blue). Bi. Waveform of mIPSPs from MOC neuron recordings (magenta line, grey shading indicates standard deviation) with an overlaid mIPSP generated from the model MOC neuron (purple). C. Waveform of control evoked PSPs from MOC neuron recordings (combined EPSP and IPSP, black line, grey shading indicates standard deviation) overlaid with PSPs from the model MOC neuron in control conditions (combined EPSP and IPSP, grey line). Ci. Waveform of excitatory evoked-PSP from MOC neuron recordings after pharmacological block of inhibitory receptors (no inhibition, EPSP only, blue line, grey shading indicates standard deviation) overlaid with PSPs from the model MOC neuron with excitatory PSPs only (no inhibition, EPSP only, dark blue line). D. Comparison of model evoked-PSPs in control (EPSP and IPSP, grey), excitatory only (inhibition removed, EPSP only, dark blue), and inhibitory only conditions (excitation removed, IPSP only, purple). E. Current-clamp recordings from MOC neurons during 100 Hz, 20 pulse MdL-stimulation to evoke APs in control (evoked EPSP and IPSP, black), and excitatory only conditions (inhibition blocked, EPSP only, blue). F. Response of the model MOC neuron to PSPs simulated at 100 Hz, 20 pulse trains with synaptic timing mimicking MdL-stimulation synaptic timing in control (EPSP and IPSP, grey), excitatory only (inhibition removed, EPSPs only, dark blue), and inhibitory only conditions (excitation removed, IPSP only, purple).

The model MOC neuron responded to synaptic inputs based on recorded mini-EPSPs (mEPSP) and mini-IPSPs (mIPSP) with closely matched amplitudes and waveforms (**Figure 7B, Bi**), confirming that the model MOC neuron responds to synaptic inputs similarly to biological MOC neurons. Next, synaptic potentials were simulated within the model MOC neuron to mimic PSPs evoked from MdL-stimulation experiments both in control conditions, and with pharmacological blockade of inhibitory synaptic inputs (**Figure 7C, Ci**). IPSPs were then added to the model to mimic MdL-stimulation conditions, with the IPSP onset being longer latency than the EPSP onset by +2.8 ms (E-I latency difference +2.8 ms), as measured in MOC neuron MdL-stimulation recordings (**Figure 7C, D**).

The responses of the model MOC neuron to repeated synaptic inputs were then tested by comparing model responses to activity recorded in MOC neurons in response to 100 Hz trains of MdL-stimulation (**Figure 6**, **Figure 7E**). Trains of MdL-PSPs were simulated within the model, first using the relative timing of excitatory and inhibitory synaptic inputs recorded during MdL-stimulation as above (E-I latency difference: +2.8 ms) and then with IPSPs absent to mimic pharmacological blockade of IPSPs in MOC neuron recordings (**Figure 7F**). Consistent with patch-clamp data, single control PSPs (EPSPs + IPSPs) did not evoke APs in the model MOC neuron. However, summation of PSPs evoked APs (latency 38.5 ms), similar to the latency to AP in MdL-stimulation experiments (49.75±29.06 ms). IPSPs were then removed from the model and summation of EPSPs initiated APs with a shorter latency (24.4 ms). This decrease in latency to APs in the absence of IPSPs is consistent with recordings from MOC neurons, and consistent with inhibition delaying APs.

We next used the MOC neuron model to determine how the timing of inhibition relative to excitation affects MOC neuron activity in the in vivo*-*like AN-stimulation configuration. In this configuration, the median latency to synaptic inhibition is closer to that of synaptic excitation compared to the greater synaptic timing separation in the MdL-stimulation configuration, and in some cells, inhibition preceded excitation (see Figure 3,4). We first generated single PSPs with varying excitatory and inhibitory synaptic timing ranging from EPSPs preceding IPSPs from 0 to 10 ms in 1 ms increments (E-I latency difference 0 to +10 ms), and IPSPs preceding EPSPs from 0 to 10 ms in 1 ms increments (E-I latency difference 0 to -10 ms). Integrated PSPs generated from these combinations had systematically varying amplitudes (**Figure 8A**). Simulating PSPs with the most extreme observed E-I latencies from AN-stimulation recordings also resulting in integrated PSPs with varying amplitudes (**Figure 8B**).

**Figure 8.**
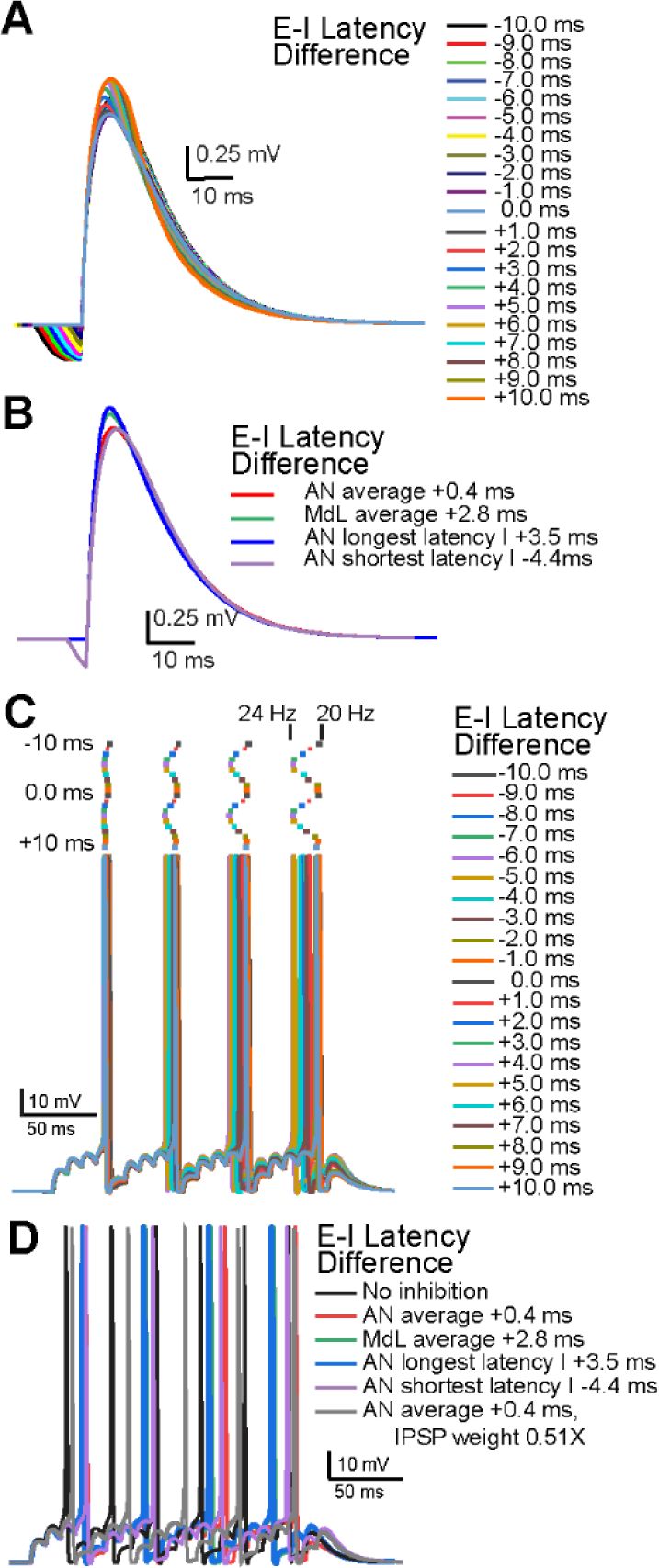
Varying latencies of summating EPSPs and IPSPs adjust model MOC neuron AP onset and rate. A. PSPs in the model MOC neuron resulting from summation of EPSPs and IPSPs at the E-I latency differences indicated. B. PSPs in the model MOC neuron resulting from summation of EPSPs and IPSPs at E-I latency differences from recorded MOC neurons. C. APs in the model MOC neuron resulting from summation of 100 Hz trains of simulated EPSPs and IPSPs at the E-I latency differences indicated. Raster plots above the APs indicate timing of AP peak. Changing E-I Latency Differences adjusted the AP rate between ∼20 and 24 Hz, indicated above the rasters. D. APs in the model MOC neuron resulting from summation of simulated EPSPs and IPSPs with E-I latency differences recorded in MOC neurons, or with reduced a IPSP amplitude (0.51 times the control value) to mimic prior synaptic depression.

We then simulated trains of PSPs in the MOC model to analyze the effect of synaptic timing on AP latency and rate. PSPs with E-I latencies from -10 to +10 ms were simulated at rates of 100 Hz. PSPs summated to evoke APs with latencies and rates that systematically varied depending on E-I latency difference (**Figure 8C**). With only EPSPs (IPSPs removed), model APs occurred at 31.5 Hz. Adding simulated inhibitory PSPs during train stimulation increased the latency to the first AP and reduced AP rate in all cases. The degree of the effect of inhibition depended on the E-I latency difference, with the most effective inhibition occurring with E-I latency differences of -10, -1, and +9 ms (AP latency: ∼48 ms, AP rate: ∼20 Hz), and the least effective inhibition occurring with E-I latency differences of -6.0 and +4.0 ms (AP latency: ∼43 ms, AP rate: ∼23.5 Hz). In wedge-slice recordings with AN-stimulation, E-I latency differences in all cells ranged from -4.4 ms to +3.5 ms, a range that nearly includes both minimal and maximal effects of inhibition on increasing AP latency and decreasing AP rate. Notably, the median AN-evoked E-I latency difference (+0.4 ms) had a greater effect on delaying and decreasing the rate of APs compared to the median MdL-evoked E-I latency difference (+2.8 ms) (**Figure 8D**), indicating that inhibitory synaptic inputs have a greater effect on suppressing APs in MOC neurons when occurring with the in vivo-like timing measured in AN-stimulation experiments. However, there was a much reduced effect of inhibition if the IPSP amplitude was decreased to match recordings in which background MNTB activity caused synaptic depression of IPSPs^19^ (**Figure 8D**). The variability in timing of inhibitory inputs recorded in different MOC neurons paired with the degree of prior synaptic depression suggests that inhibition may have a different effect on each MOC neuron in the brain, and can thus flexibly adjust MOC neuron activity and de-synchronize the population of MOC neurons.

Together, our results indicate that inhibition from the MNTB can delay initial spiking of MOC neurons, and that the precise timing of inhibition can flexibly adjust MOC activity rates. In addition, the inhibitory pathway to MOC neurons is exceptionally fast and precise, due to the properties of synaptic transmission and neuron function localized to the AN synapses onto CN cells, the CN cells themselves, and CN projection axons.

## Discussion

### MOC neuron activity

Diverse synaptic inputs to MOC neurons likely converge to adjust MOC function under changing hearing conditions. The sound responses of MOC neurons in vivo have low thresholds, V-shaped tuning curves, sound-evoked firing rates up to 110 Hz, and a variable latency from sound onset to APs that decreases with loud sounds^24,25,48–50^. Histological and lesion experiments have shown synaptic inputs to MOC neurons from a variety of auditory and non-auditory neurons^36,38,39,51–62^. In vitro patch-clamp recordings of synaptic inputs to MOC neurons include functional demonstration of ascending excitatory synaptic inputs from CN T-stellate cells^17^ and possibly CN SCC neurons^37^, as well as descending, facilitating excitatory synaptic inputs from the IC^17^. Inhibitory synaptic inputs were recorded in putative MOC neurons^63^ and more recently originating in the MNTB as shown in genetically identified MOC neurons^18^. Serotonergic excitation of MOC neurons^64^ suggests function of additional modulatory inputs. These diverse inputs likely underestimate the convergence of synaptic inputs to MOC neurons. The in vitro wedge-slice preparation combined with machine learning and single-neuron computational modeling used here is a first step in probing synaptic integration in MOC neurons in vitro, from monaural ascending circuits with in vivo-like timing.

### Functional synaptic inputs to MOC neurons

The wedge-slice includes the monaural ascending circuitry to MOC neurons beginning with AN axon terminals from Type I SGN onto CN neurons including GBC, T-stellate cells, and other CN cells. CN axons projecting to the SOC are intact^32^. Type II SGN terminations in the CN including onto granule and other cells^65–67^ are also present. Intrinsic CN circuitry remains intact, including excitatory and inhibitory connections within the CN^68–80^ and reciprocal connectivity between bushy^81^ and T-stellate cells^82^. Absent from the preparation are the cochlea, commissural pathways between the CN, and most descending circuits to the CN. While we did not determine the cell types generating synaptic responses in MOC neurons, the varied patterns of multiple peaks of PSCs, found both in AN- and MdL-stimulation experiments, suggest diverse excitatory and inhibitory inputs. These patterns suggest either additional direct CN projections, or poly-synaptic inputs with a contralateral CN origin. Additional potential sources of excitation are SCC neurons^37^, axon collaterals from the GBCs^83,84^, or other as-yet unidentified sources. For inhibition, there may be an additional direct projection from the CN^85^, but poly-synaptic inputs are likely because of the ∼3-4 ms delay between first and second clusters of IPSCs in most recordings. Potential poly-synaptic inhibitory inputs from auditory neurons include the LNTB^86,87^, the superior peri-olivary nucleus (SPON)^88,89^, other inhibitory neurons in the VNTB^90^, or the MOC neurons themselves, which are cholinergic but likely also GABAergic^91–93^.

### Speed and fidelity of the GBC–MNTB inhibitory pathway

The GBC-MNTB pathway inhibits many other SOC neurons, including the LSO, MSO, lateral olivocochlear neurons (LOC), and SPON neurons. The pathway has large Endbulb of Held synapses from the AN onto the GBCs, while the GBCs have large diameter, heavily myelinated axons^29,94–96^, and a termination in the large Calyx of Held onto MNTB neurons. High fidelity synaptic transfer throughout the pathway allows activity up to hundreds of Hz^94,97–100,101–113^. The high speed of this pathway was demonstrated in the GBC–MNTB synapses to the MSO^28^ in gerbils. In our present results in mice, in two MOC neuron recordings the inhibitory pathway is shorter latency than the excitatory pathway, despite having an additional synapse, indicating that despite lacking some of the axonal GBC specializations that are present in gerbils^31^, mice also have an exceptionally fast GBC-MNTB pathway. However, in our results the excitatory pathway on average has a shorter latency than the inhibitory pathway. One interpretation of this is that in addition to the inhibitory pathway being fast, the excitatory pathway is also exceptionally fast, which is unexpected due to the ∼5 ms AP latency in T-stellate cells^114^. Another possibility is that MNTB axon branches to MOC neurons are slower conducting compared to MNTB axon branches to MSO neurons.

Our results also pinpoint the segments of the inhibitory pathway responsible for both its speed and precision by directly comparing the relative latency differences, jitter, and response probability between EPSCs and IPSCs evoked using MdL- vs AN-stimulation. Studies have indicated that, although GBC’s have large^33,34,43,115^, often supra-threshold^45,70,115,116^ PSPs, some amount of synaptic summation is required for the precise phase-locking and enhanced entrainment of GBC’s^33–35,114^ that is also attributed to a low somatic resistance with a short temporal integration window^43,45,117^. In our results, addition of the CN with AN-stimulation reduces the probability of activity in the pathway, confirming that there is not a 1:1 PSP:AP relationship and that synaptic summation contributes to the ‘coincidence detector’ function of GBC’s. There may also be species-specific circuit differences, intact inhibitory CN circuits, decreased release probability from AN axons, or smaller post-synaptic responses in the GBC. Our observed decreased latency of inhibitory PSCs relative to excitatory PSCs was also only apparent when the circuit included the full CN via AN-stimulation, not with MdL-stimulation that lacked GBC participation, pinpointing the speed of the inhibitory pathway to the CN and GBCs. Finally, with MdL-stimulation, the inhibitory pathway has more synaptic jitter than the excitatory pathway. However, this is compensated for in the AN-stimulation experiments to result in equal total jitter between the excitatory and inhibitory pathways. In accordance with the precision of the endbulb of Held– GBC synapse^72^, this suggests that this portion of the inhibitory pathway that is added during AN-stimulation experiments relative to MdL-stimulation experiments is so precise that it can compensate for increased jitter at later synapses.

### Integration of excitation and inhibition at MOC neurons

Our findings from both AN-stimulation experiments in wedge-slices and the computational MOC model inform knowledge of the role of integration of synaptic inhibition and excitation on MOC neurons both at the single cell and the population level. The presence of any inhibition in the model reduced AP rates from control values. In recordings, inhibition was variable and sometimes shorter latency relative to excitation. Then, in the MOC neuron computational model, changing synaptic timing to reflect the different latencies of EPSCs relative to IPSCs (E-I latency difference) resulted in variable amplitude summed PSPs and variable AP rates. E-I latency differences that corresponded to values from MOC neuron recordings ranged from having a minimal effect on reducing AP rates when inhibition lags excitation, to having the maximal effect when inhibition slightly precedes excitation by ∼1 ms. Interestingly, the most striking example in which inhibition preceded excitation by 4.4 ms did not have the strongest effect on MOC AP rates in the model, suggesting that closely-timed synaptic inhibition and excitation are most effective at reducing MOC neuron activity. Further, we speculate that if the timing of inhibition is different across the population of MOC neurons, in addition to the more variable excitatory timing, each cell will have slightly different AP latencies and rates in response to a sound, thus de-synchronizing MOC activity across the population.

Our results present a model that while MOC neurons receive rapid and precise excitation at the single cell level, multiple mechanisms are in place to syncopate, broaden, and smooth activity at the population level. This extends from broadly tuned inhibitory inputs^18^, variable axonal projection patterns in the cochlea ^24,25,118–122^, low synaptic release probability paired with synaptic facilitation at MOC axon terminals onto cochlear outer hair cells (OHC)^123^, and a slow post-synaptic response in the OHC. Our results add fast and precisely timed synaptic inhibition to the list of mechanisms to smooth MOC efferent activity, which paradoxically adds imprecision to the MOC system to exert slow effects on cochlear gain control.

### Methods Summary

Asymmetric ‘wedge slice’ brain slices were used to perform patch-clamp recordings from identified MOC neurons in ChAT-IRES-Cre; tdTomato P13-21 mice. Electrical stimulation of pre-synaptic axons either at the midline (MdL) or at the intact auditory nerve (AN) root projecting into the contralateral cochlear nucleus (CN) evoked multicomponent post-synaptic currents (PSCs) consisting of both excitatory and inhibitory inputs. During current-clamp recordings, post-synaptic potentials (PSP) were evoked by MdL stimulation of pre-synaptic axons in the presence and absence of blockers of inhibitory neurotransmission (50 µM gabazine, 1 µM strychnine), to test the effect of inhibition on PSP summation. Calcium imaging in wedge slices of Atoh7/Math5 Cre; GCaMP6f during AN stimulation was performed with and without blockers of glutamatergic receptors (5 µM CNQX, 50 µM APV) to ensure that AN stimulation acted via synaptic, not direct electrical, excitation of CN neurons. A machine learning RandomForest paradigm was trained to classify EPSCs and IPSCs recorded in MdL stimulation experiments for which excitatory vs inhibitory PSC identity was known, then the trained machine learning paradigm was used to classify PSCs from AN experiments for which excitatory vs inhibitory identity was unknown. A computational model of a single MOC neuron was designed based on properties recorded from MOC neurons, then synaptic inputs were simulated within the model to reflect the varying latencies of excitatory and inhibitory synaptic inputs recorded in MOC neurons with MdL vs AN stimulation.

## Acknowledgements

Thank you to Dr. Lena Ebbers and Dr. Hans Nothwang from Carl von Ossietzky Universität Oldenburg for the Atoh7/Math5 Cre mice. Thank you to Dr. Kirupa Suthakar for thoughtful discussions and a critical reading of the manuscript. This research was supported by the Intramural Research Program of the NIH, NIDCD, Z01 DC000091 (CJCW) and the Bioinformatics and Biostatistics Core of the NIH / NIDCD.

## Supplementary Methods

### RESOURCE AVAILABILITY

#### Materials availability

This study did not generate new unique materials.

#### Data and code availability

##### DATA

Electrophysiology and imaging data will be deposited and publicly available by the date of publication.

##### CODE

All original code will be deposited and will be publicly available as of the date of publication.

##### ANALYSIS TOOLS

Any additional information required to reanalyze the data reported in this paper is available from the lead contact upon request.

## EXPERIMENTAL MODEL DETAILS

### Ethical approval and animal housing

Animal procedures followed National Institutes of Health guidelines, as approved by the National Institute of Neurological Disorders and Stroke/National Institute on Deafness and Other Communication Disorders Animal Care and Use Committee. Pre-weaned mice postnatal age 13-21 (P13-P21) were used for experiments and were housed with parents and littermates before use. Mice were housed in the NIDCD animal facility with a 12:12 light/dark cycle where food and water were provided *ad libitum*. Mice of both sexes were used for experiments. For consideration of sex as a biological factor, postsynaptic currents (PSCs) recorded during stimulation of the ventral acoustic stria (‘midline stimulation’; MdL-stimulation) were analyzed and compared between the sexes (see below for complete methods). No significant differences were found for excitatory or inhibitory PSCs using the metrics of onset latency, onset jitter, rise time, decay tau, amplitude and probability. Datasets were therefore pooled.

**Table M1:** Values for metrics analyzed for PSCs evoked with midline (MdL) stimulation for comparison of sex as a biological factor. n # for each group in parentheses represents # of peaks (clusters) analyzed. Comparisons made between sexes showed no significant differences with Mann-Whitney U Test.

| Median ± MAD | | Onset<br>Latency (ms) | Onset<br>Jitter (ms) | Rise<br>Time (ms) | Decay<br>Tau (ms) | Amplitude<br>(nA) | Probability |
| --- | --- | --- | --- | --- | --- | --- | --- |
| Male | EPSCs (10) | 2.11 ± 0.43 | 0.19 ± 0.08 | 0.61 ± 0.28 | 2.44 ± 0.50 | -0.05 ± 0.02 | 0.48 ± 0.21 |
| Female | EPSCs (11) | 2.02 ± 0.67 | 0.41 ± 0.15 | 0.73 ± 0.20 | 2.23 ± 0.55 | -0.04 ± 0.02 | 0.52 ± 0.31 |
| Male | IPSCs (8) | 5.01 ± 1.98 | 0.53 ± 0.08 | 1.26 ± 0.59 | 5.79 ± 2.85 | -0.07 ± 0.02 | 0.39 ± 0.11 |
| Female | IPSCs (10) | 5.30 ± 1.26 | 0.44 ± 0.20 | 1.40 ± 0.32 | 3.34 ± 0.48 | -0.05 ± 0.02 | 0.25 ± 0.16 |

### ChAT-IRES-Cre x tdTomato mouse line

ChAT-IRES-Cre transgenic mice on either a C57BL/6J (RRID:IMSR_JAX:028861) or a C57BL/6N (RRID:IMSR_JAX:018957) background strain were crossed with tdTomato reporter mice (Ai14, Cre reporter allele inserted into Rosa 26 locus; RRID:IMSR_JAX:007914) to yield offspring heterozygous for each allele for experiments. These mice were used to target MOC neurons for patch-clamp recordings as previously described (Torres Cadenas et al., 2020).

### Atoh7/Math5 Cre x GCaMP6f mouse line

Atoh7/Math5 Cre mice (Yang et al., 2003, RRID:MGI:3717726) were crossed with GCaMP6f Ai95(RCL-GCaMP6f, RRID:IMSR_JAX:028865) mice for use in calcium imaging experiments of cochlear nucleus bushy cells.

### Brain slice preparation

Mice were killed by carbon dioxide inhalation at a rate of 20-30% of chamber volume per minute, then decapitated. The brain was removed in cold artificial cerebrospinal fluid (aCSF) containing the following (in mM): 124 NaCl, 1.2 CaCl_2_, 1.3 MgSO_4_, 5 KCl, 26 NaHCO_3_, 1.25 KH_2_PO_4_, and 10 dextrose; 1 mM kynurenic acid was included during slice preparation. The pH was equal to 7.4 when bubbled with 95% O_2_/5% CO_2_. In experiments in which mini-post-synaptic potentials (mPSPs) were recorded, 1 µM tetrodotoxin (TTX) was included in the aCSF. Asymmetric slices were obtained as previously described (Fischl & Weisz 2020). Briefly, the brainstem was carefully dissected from the skull to maintain a portion of the auditory nerve roots entering the cochlear nucleus, similar to methods used for thick slice preparations ^27,28^. Then, a wedge-shaped section was acquired using a stage with an adjustable angle such that the lateral edge of one hemisphere was approximately 1-1.2 mm thick and contained the cochlear nucleus, and the opposite lateral edge was ∼200 μm, creating a thickness of ∼300-400 μm where MOC neurons were identified for patch-clamp experiments in the ventral nucleus of the trapezoid body (VNTB). An additional 300 μm slice was obtained rostral to the ‘wedge slice’ and used for additional experiments. For some experiments utilizing midline stimulation for evoked PSPs or for recordings of miniPSPs, symmetrical brain slices were prepared as previously described ^18^.

Sections were transferred to an incubation chamber and maintained at 35 ± 1°C for 30-60 min. Slices then cooled to room temperature until used for experiments within 4 hours of slicing. Wedge slices were used immediately after a short recovery incubation (30 min) to improve cell viability which tends to diminish more rapidly than typical symmetrical slices due to reduced aCSF solution penetration in the larger tissue volume.

### Patch-clamp recordings

Sections were transferred to a recording chamber which was continuously perfused with aCSF at a rate of 5-10 mL/min. The bath temperature was held at 35 ± 1°C using an in-line heater (Warner) coupled to a temperature controller (Warner). The tissue was viewed using a Nikon Eclipse Ni-E microscope with an Apo LWD 25X/1.10 NA water-immersion objective attached to a Retiga Electro CCD camera (QImaging) operated using NIS Elements software (version 4.51.01). Epifluorescence illumination with red emission filters were used to locate MOC neurons in the VNTB for recordings. Targeted cells were then observed using DIC optics for patch-clamp recordings.

Pipettes for patch-clamp recordings were pulled from 1.5 mm borosilicate glass capillaries to resistances between 3-7 MOhm. For voltage-clamp experiments an internal solution containing (in mM) 76 Cs-methanesulfonate, 56 CsCl, 1 MgCl_2_, 1 CaCl_2_, 10 HEPES, 10 EGTA, 0.3 Na-GTP, 2 Mg-ATP, 5 Na_2_-phosphocreatine, 5 QX-314, and 0.01 Alexa Fluor-488 hydrazide was used. The pH was adjusted to 7.2 with CsOH. For current-clamp experiments, an internal solution containing (in mM) 125 K-gluconate, 5 KCl, 1 MgCl_2_, 0.1 CaCl_2_, 10 HEPES, 1 EGTA, 0.3 Na-GTP, 2 Mg-ATP, 1 Na_2_-phosphocreatine, and 0.01 Alexa Fluor-488 hydrazide was used. The pH was adjusted 7.2 with 1N KOH. Liquid junction potentials were -6 mV, CsCl solution and -2 mV, KGlu solution and were not adjusted for. Electrophysiology recordings were performed using a HEKA EPC10 amplifier controlled using PatchMaster NEXT (version 1.1). The recordings were sampled at 50 kHz and filtered on-line at 10 kHz. Series resistance was compensated between 60-85%. Cells were voltage-clamped at -60 mV unless stated otherwise. In current-clamp, holding currents were injected to maintain the baseline membrane potential at -60 mV.

### Stimulation of auditory nerve and ventral acoustic stria

Post-synaptic currents (PSC) and post-synaptic potentials (PSP) recorded in MOC neurons were evoked by electrical stimulation of axons using a bipolar tungsten electrode (WPI). For auditory nerve (AN) stimulation, the electrode was lowered onto the approximate center of the auditory nerve root diameter between the cut end of the nerve and its entry into the CN. For stimulation at the midline, the electrode was placed just lateral to the midline (contralateral hemisphere to the MOC neuron recording) near the ventral surface of the tissue onto fibers of the ventral acoustic stria. The stimulation was applied with an Iso-Flex Stimulus Isolation Unit (A.M.P.I.) and the intensity was adjusted to obtain consistent amplitude postsynaptic responses in MOC neurons (stimulation range 10-2000 μA). In two experiments using AN-stimulation, the stimulus intensity was increased until the PSC latencies jumped to a shorter value (see figure 3,4), indicating direct recruitment of CN axons and bypassing of auditory nerve synapses onto CN cells. To isolate inhibitory currents in voltage-clamp, the membrane potential was clamped at 0 mV, the approximate reversal potential for AMPA-mediated glutamatergic currents. Inhibitory inputs were blocked where indicated with bath application of strychnine (1 µM) and Gabazine (SR95531, 50 µM).

### Miniature PSP recordings

For miniature PSP (mPSP) recordings, 1 µM tetrodotoxin (TTX, Alomone Labs) was included in the aCSF. Recordings were performed for several minutes to collect baseline (control) mPSPs. Inhibitory inputs were blocked using bath application of strychnine (1 µM) and Gabazine (SR95531, 50 µM) and gap free recordings were again taken during pharmacological manipulation. mPSPs were detected using using MiniAnalysis software version 6.0.7 (Synaptosoft) using a threshold of 2 x RMS noise. PSCs were then accepted or rejected based on the characteristic PSC waveform. The decay time constants of PSCs were calculated in Mini Analysis software from individual events.

### Calcium Imaging

Calcium imaging of activity in CN bushy cells was performed using the Atoh7/Math5 Cre mouse line crossed with a GCaMP6f mouse line (see above). Asymmetric wedge slices were prepared as described above (Experimental Model Details). aCSF used for recording calcium signals was modified to contain 2 mM CaCl_2_. Epifluorescence illumination with green emission filters were used to locate bushy cell neurons in the anteroventral cochlear nucleus (AVCN). The AVCN was targeted to increase the potential for imaging of globular bushy cells (GBCs) which project to the contralateral MNTB. Calcium signals were imaged using a Nikon Eclipse Ni-E microscope with an Apo LWD 25X/1.10 NA water-immersion objective in 2-photon excitation mode at 920 nm (Mai Tai HP, Spectra-Physics). A single focal plane was imaged for data collection. Imaging was performed at 3 Hz for 20 seconds using the resonant scanning galvos (Nikon Elements software version 4.51.01). Protocols consisted of 5 seconds of baseline data collection followed by auditory nerve stimulation in 3 bouts (each bout is 20 pulses at 100 Hz), at 5 seconds intervals, followed by an additional 5 seconds after the third stimulation bout, then at least 30 seconds without imaging or stimulation. Control data was acquired for several minutes before glutamate receptor blockers were applied. In a subset of experiments, CNQX (5 μM) was applied alone to block ionotropic glutamate receptors. In remaining calcium imaging experiments, APV (50 μM) was also added to additionally block NMDA receptors. Acquisition protocols were repeated during blocker application. Blockers were applied for ∼10 minutes before washout. After washout, protocols were repeated to assess recovery of control conditions.

### Computational MOC neuron model

A model of a single MOC neuron was constructed using NEURON v8 ^124^ and Python 3. The neuron topology was generated from a published MOC neuron morphology (Figure 9 from, ^47^). The number of segments was empirically determined to be 86, inserted using the nseg function, and compartments were organized into larger morphological groups including the soma (length 33.6 µm, diameter 6.1 µm), axon (length 180.0 µm, diameter 1 µm), and dendrites. The three primary dendrites projected from the soma (lateral: length 12.8 µm, diameter 1 µm), dorsal (length 10.1 µm, diameter 1 µm), and medial (length 7.3 µm, diameter 1 µm). Each of these three primary dendrites branched further into 8 to 46 dendrites with lengths from 1.5 to 31.4 µm, and diameters from 1-2 µm. This detailed neuron topology allowed specific control of physiological properties. The uniform axial resistance was 210 ohm-cm, and membrane capacitance was 1 µF/cm^2. Channels were inserted into the membrane (Table 1) to recapitulate our experimental results. HCN channel reversal potential was set to -38 mV. The model was run at 35 C.

Post-synaptic potentials (PSP) were simulated at the soma to replicate recorded values. The model MOC neuron responded to synaptic inputs based on recorded mini-EPSPs (mEPSP) and mini-IPSPs (mIPSP) with output amplitudes and waveforms closely matching recorded values (mean±SD; mEPSP: amplitude: 0.86±0.23 mV, time constant of decay: 8.65±1.43 ms, n=6 neurons; mIPSP: amplitude: -1.57±1.41 mV, time constant of decay: 18.1±9.68 ms, n=7 neurons; model MOC mEPSP: amplitude: 0.90 mV, time constant of decay: 8.21 ms; model MOC mIPSP: amplitude: -1.56 mV, time constant of decay: 14.4 ms.

Next, synaptic potentials were simulated within the model MOC neuron to mimic PSPs evoked from MdL-stimulation experiments both in control conditions, and with pharmacological blockade of inhibitory synaptic inputs (MOC recording control evoked-PSP: amplitude: 1.88±0.95 mV, time constant of decay: 10.7±4.00 ms, n=9 neurons; inhibition blocked evoked-PSP (EPSP only): amplitude: 2.20±1.17 mV, time constant of decay: 14.1±7.50 ms, n=9 neurons; model MOC control evoked-PSP: amplitude: 1.64 mV, time constant of decay: 14.8 ms; model MOC inhibition blocked evoked-PSP (EPSP only): amplitude: 1.91 mV, time constant of decay: 16.4 ms. To achieve these output parameters, model MOC neuron input values were as follows: MOC model EPSP input parameters: rise time = 0.2 ms, time constant of decay = 6 ms, synaptic weight = 0.0015, reversal potential = 0 mV, resulting amplitude = 8.506 mV). The inhibitory PSP was designed so that simulation of EPSPs and IPSPs in the model replicated control MOC neuron current-clamp recordings of evoked PSPs. The model MOC IPSP output parameters were amplitude: -0.269 mV, time constant of decay: 14.8 ms. To achieve these values, the model MOC IPSP input parameters were as follows: rise time = 2.75 ms, time constant of decay = 3.64, synaptic weight 0.00032, reversal potential = -90 mV, resulting amplitude = -0.536 mV).

PSPs with excitatory and inhibitory components were simulated with onset timing that systematically changed in 1 ms increments from excitation-inhibition latencies (E-I latency) from -10 (IPSPs precede EPSPs by 10 ms) to +10 (EPSPs precede IPSPs by 10 ms). E-I latencies were also simulated within the model to replicate recorded PSC latencies (Results text). PSPs were then simulated within the model in trains of 20 pulses at 100 Hz, using either the excitation-only PSP, or the combined PSPs with E-I latencies that systematically varied between -10 and +10, in 1 ms intervals.

The model will be available at Model dB (identifier # TBD at acceptance)

**Methods Table 1.** Biophysical properties of a modeled MOC neuron compared to measured values recorded from MOC neurons.

| Channels | Location (segments) | Conductance |
| --- | --- | --- |
| High Threshold $K^+$ (h.HT) | All | 0.02 S/cm<sup>2</sup> |
| Low Threshold $K^+$ (h.LT) | All | 12 mS/cm<sup>2</sup> |
| Low Threshold $K^+$ (h.kbl LT) | All | 0.6 ms<sup>-1</sup> |
| H-H type $Na^+$ (h.na) | All | 8 nS/cm<sup>2</sup> |
| HCN (h.Ih 400t8) | All | 12 μS/cm<sup>2</sup> |
| High Threshold $K^+$ (h.HT) | Axon | 0.32 mS/cm<sup>2</sup> |
| Low Threshold $K^+$ (h.LT) | Axon | 6 cS/cm<sup>2</sup> |
| H-H type $Na^+$ (h.na) | Axon | 9 kS/cm<sup>2</sup> |

## QUANTIFICATION AND STATISTICAL ANALYSIS

### Statistics for PSCs

Analysis of synaptic inputs to MOC neurons required classification of evoked PSCs as excitatory or inhibitory. Clampfit software was used to detect individual PSCs for analysis. Latency to PSC onset, rise time, latency to PSC peak, amplitude and decay time constant (tau decay) were measured for PSCs recorded at both –60 mV and 0 mV holding potential. A single exponential function fit was used to calculate the time constant of decay (τ). These data were then used for individual cell clustering analysis (below).

### Individual cell clustering analysis

Clustering analysis was performed using PSC metrics in order to sort PSCs into statistically defined clusters. Clustering was performed with PSCs collected at both –60 mV and 0 mV holding potential, when available. For each cell’s PSCs, values for onset latency, rise time, amplitude and decay time constant collected from Clampfit were imported into R. Function libraries utilized for clustering included {parameters}, {factoextra} and {NbClust}. The appropriate number of clusters was determined using the gap statistic method (Tibshirani et al. 2001). This method was chosen because of its ability to select “one” as the optimal number of clusters where appropriate. The “clusGap” function was used with kmax=10 (max# of clusters), nstart=25 (# of random start centers) and B=500 (bootstrapping). Once the appropriate number of clusters was determined, a k-means cluster analysis was performed in R using the “kmeans” function to sort the PSCs into clusters (with centers=output # from gap statistic analysis and nstart=25). Once the PSCs were assigned a cluster number, statistical analyses were performed on each cluster. For PSCs collected with midline stimulation where data was acquired at 0 mV holding potential, a cluster was deemed inhibitory if it was present at both -60 mV and 0 mV. A cluster was categorized as excitatory if PSCs from the cluster were only present at -60 mV. This was a robust categorization, with only two out of eleven cells having a PSC mis-identified in the cluster, in both cells a single excitatory PSC was classified into an inhibitory cluster (2 out of 654 mis-identified PSCs). PSCs acquired with auditory nerve (AN) stimulation were analyzed via the same cluster analysis. AN stimulation PSCs were categorized as excitatory or inhibitory using a machine learning algorithm (see below). Statistical comparisons between excitatory and inhibitory PSC clusters were then made across the population.

### Machine Learning Algorithm to classify post-synaptic currents (PSCs)

A RandomForest machine learning algorithm was utilized to classify recorded PSCs as excitatory or inhibitory based on the variables of PSC rise time, time constant of decay, amplitude, probability of occurrence within a cluster, onset jitter within a cluster, peak jitter within a cluster, and animal age. All values were continuous except for animal age, which was treated as a categorical variable. The model included 952 PSCs recorded in the ML-stimulation configuration at a holding potential of -60 mV, from 22 MOC neurons. These PSCs had been previously classified as excitatory or inhibitory based on recordings from the same neurons at a holding potential of 0 mV and the clustering analysis described below. After training the RandomForest algorithm on this data, the model classification accuracy reached 99.89%, with out-of-bag (OOB) error stabilization at 150 ‘trees’. Re-running the algorithm on the training dataset determined that it was able to distinguish excitatory and inhibitory events perfectly with an area under the curve (AUC) of 1, indicating excellent ability of the model to distinguish excitatory vs inhibitory PSCs. The model was then used to classify the individual PSCs in the AN stimulation dataset recorded at -60 mV as excitatory or inhibitory. First, the RandomForest algorithm determined the probability that each of the 344 PSCs was excitatory or inhibitory. PSCs were given the classification that had the highest probability (>0.5) by the algorithm. The algorithm gave slightly higher classification probabilities for excitatory (0.81±0.14, n=181) compared to inhibitory (0.75±0.12, n=163) PSCs. AN stimulation PSCs were grouped into 28 clusters determined above through cluster analysis. Three clusters contained both excitatory and inhibitory PSCs. Two cells had a majority of one classification (7 of 10 excitatory and 19 of 21 excitatory and one cell was approximately split 6 of 11 excitatory). For further analyses, we separated these three mixed clusters into an excitatory and inhibitory cluster each to yield a total of 31 AN-stimulation clusters.

### Calcium Imaging

After acquisition of fluorescent signals in cochlear nucleus bushy cells in Atoh7/Math5 Cre; GCaMP6f wedge slices (see above), fluorescence changes in response to electrical stimulation of axons were measured to determine the effect of synaptic stimulation of bushy cells with and without blockers of post-synaptic receptors. Polygonal ROIs were drawn by hand around neurons and any major processes that could be resolved (Elements software version 4.51.01), and average intensity values for ROIs were calculated for each frame. Maximum fluorescence elicited from AN stimulation within a protocol was compared to baseline average (F) of each ROI. Fluorescence change (ΔF) and relative fluorescence change (ΔF/F) was calculated using Excel. Heat maps were constructed from the intensity value output of a given frame from the Elements software using custom MATLAB scripts. Baseline values were calculated as the average pixel intensity of the first 15 frames (∼5 seconds) before axon stimulation. Cells were considered active if the average fluorescence of an ROI reached two standard deviations (SD) above the baseline average in at least 2 of the 3 stimulations during a protocol. Active cells were then used to compare the ΔF/F between control and glutamate block conditions.

### Data analysis and statistics

Statistical analyses were performed in Origin (v2021 and v2022). Normality tests were performed on data sets using the Shapiro-Wilk test. The majority of data sets were non-normally distributed so non-parametric testing was employed. The Mann-Whitney U test was used for testing between two independent groups. A One-Sample Wilcoxon Signed Rank Test determined whether E-I latency difference values were significantly different from zero for MdL- and AN-stimulation PSCs. Action potential metrics collected at different stimulus rates (Figure 6) were compared in the control condition using Friedman’s ANOVA. Post-hoc Dunn’s test was used to test significance between stimulus rates. Action potential metrics were also compared between control and inhibition block conditions using paired Wilcoxon Signed Rank Test. Calcium imaging data were analyzed using Kruskal-Wallis ANOVA where post-hoc Dunn’s test was used to test whether control data sets between the CNQX and CNQX+APV conditions were significantly different from each other. Population data are summarized in box plots with the box representing the 1^st^ and 3^rd^ quartiles, the line representing the median, the square representing the mean and the error bars representing the 10^th^ and 90^th^ percentiles. Figures were prepared in Origin and Adobe Illustrator.

## Preprint Server

This manuscript was submitted to BioRXiv: https://doi.org/10.1101/2023.12.21.572886. The copyright holder for this preprint is the author/funder, who has granted bioRxiv a license to display the preprint in perpetuity. This article is a US Government work. It is not subject to copyright under 17 USC 105 and is also made available for use under a CC0 license.

## Classification

Biological Sciences, Neuroscience

## Supplementary Tables

**Table S1.** Parameters of MdL-stimulation-evoked PSCs from Figure 1D, for all clusters.

|  | MdL-EPSC | MdL-EPSC n<br>(clusters / cells) | MdL-IPSC | MdL-IPSC n<br>(clusters / cells) | p-value<br>(Mann-Whitney U Test) |
| --- | --- | --- | --- | --- | --- |
| Amplitude, -60 mV (pA) | 66.95±3.95 | 25/17 | 55.39±3.10 | 21/15 | 0.69 |
| Rise Time (ms) | 0.63±0.19 | 25/17 | 0.86±0.15 | 21/15 | 8.02E-04 |
| Time constant of decay ($\tau$, ms) | 2.23±0.52 | 23/17 | 3.51±0.93 | 21/15 | 4.95E-07 |
| Latency (ms) | 1.92±0.37 | 18/18 | 4.47±0.93 | 15/15 | 4.79E-06 |

**Table S2.** Top: Fluorescence changes in Atoh7/Math5Cre; GCaMP6f expressing bushy cells in control conditions, with CNQX, and wash of CNQX. Bottom: Fluorescence changes in control, with CNQX + APV, and wash of CNQX + APV. Differences between conditions tested with Friedman’s ANOVA followed by post-hoc Dunn’s Test. % suppression from control: (control DF/F-block DF/F) / (control DF/F-1)*100%. Number of cells for each set of experiments in parentheses.

| | $\Delta F/F$ | p-value | % suppression from control |
| --- | --- | --- | --- |
| control (106) | 1.176±0.075 |  |  |
| CNQX | 1.059±0.018 | 1.04E-20 | 67.67±12.43 |
| CNQX wash | 1.150±0.073 | 2.74E-13 |  |
| control (121) | 1.160±0.071 |  |  |
| CNQX + APV | 1.039±0.009 | 1.14E-34 | 74.51±11.31 |
| CNQX + APV wash | 1.125±0.055 | 2.54E-24 |  |

**Table S3.** Parameters of AN-stimulation evoked PSCs from Figure 4E, for all clusters.

|  | AN-EPSC | AN-EPSC n<br>(clusters / cells) | AN-IPSC | AN-IPSC n<br>(clusters / cells) | p-value<br>(Mann-Whitney U Test) |
| --- | --- | --- | --- | --- | --- |
| Amplitude -60 mV (pA) | 60.6±39.0 | 13/8 | 25.6±11.1 | 17/10 | 0.14 |
| Rise Time (ms) | 0.86±0.15 | 13/8 | 0.92±0.31 | 17/10 | 0.35 |
| Time constant of decay ($\tau$, ms) | 2.80±1.65 | 11/8 | 3.59±1.50 | 16/10 | 0.67 |
| Latency (ms) | 5.83±1.22 | 13/8 | 6.71±0.87 | 17/10 | 0.04 |

**Table S4.**
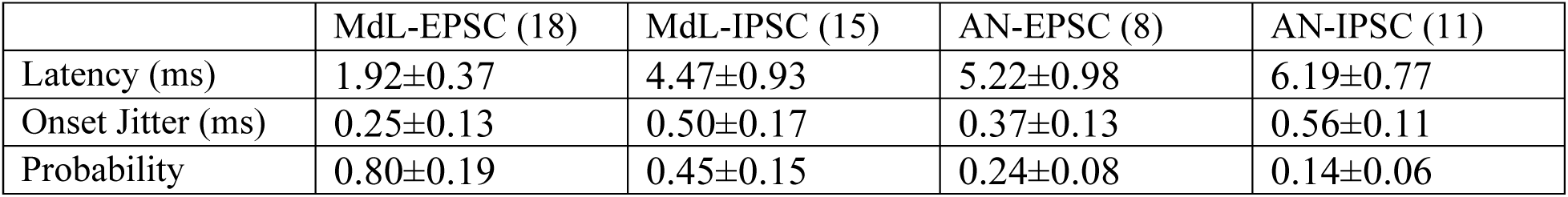
Parameters of the first cluster of PSCs evoked from MdL- and AN-stimulation from Figure 5. Numbers in parentheses indicate number of cells.

**Table S5.** Measurements of AP probability, rate, the number of stimulations to first AP, and latency to first AP in 5-8 MOC neuron recordings in response to MdL-stimulation at the frequency indicated, from Figure 6C. Measurements are presented in control, during pharmacological blockade of post-synaptic inhibitory receptors (“block”), and wash conditions, as median±MAD. Numbers in parentheses indicate n. # indicates significantly different from control 10 Hz stimulation (Friedman ANOVA with post-hoc Dunn’s test). * indicates significantly different from control within stimulus frequency comparison (P<0.05, Paired Wilcoxon Signed-Rank Test).

| Rate | Treatment (n) | AP Prob | AP Rate (Hz) | # stimulations to first AP | Latency to first AP |
| --- | --- | --- | --- | --- | --- |
| 10 Hz | Control (7) | 0 | 0 | >20 | >2000 |
| 10 Hz | Block (7) | 0 | 0 | >20 | >2000 |
| 10 Hz | Wash (5) | 0 | 0 | >20 | >2000 |
| 50 Hz | Control (7) | 0.005±0.005 | 0.25±0.25 | 19.0±1.0 | 379.2±20.9 |
| 50 Hz | Block (7) | 0.053±0.053 | 2.63±2.63 | 7.80±4.40 | 144.4±86.87 |
| 50 Hz | Wash (5) | 0 | 0 | >20.0 | >400 |
| 100 Hz | Control (7) | 0.035±0.015 | 3.49±1.51 | 14.15±4.30 | 139±44.6 # |
| 100 Hz | Block (7) | 0.078±0.036 * | 7.78±3.75 * | 5.40±2.76 * | 49.8±29.1 * |
| 100 Hz | Wash (5) | 0.051±0.03 | 5.08±3.00 | 9.50±5.13 | 126±56.7 |
| 200 Hz | Control (7) | 0.045±0.040 # | 9.00±8.00 # | 14.2±4.20 # | 69.3±22.3 # |
| 200 Hz | Block (7) | 0.075±0.047 * | 15.0±9.40 * | 6.00±2.00 * | 26.3±12.0 * |
| 200 Hz | Wash (5) | 0.013±0.013 | 2.67±2.67 | 16.6±3.50 | 75.7±19.5 |

